# The apical region of the herpes simplex virus major capsid protein promotes capsid maturation

**DOI:** 10.1101/319988

**Authors:** Laura L. Ruhge, Alexis G. E. Huet, James F. Conway, Gregory A. Smith

## Abstract

The herpesvirus capsid assembles in the nucleus as an immature procapsid precursor built around viral scaffold proteins. The event that initiates procapsid maturation is unknown, but it is dependent upon activation of the VP24 internal protease. Scaffold cleavage triggers angularization of the shell and its decoration with the VP26 and pUL25 capsid-surface proteins. In both the procapsid and mature angularized capsid, the apical region of the major capsid protein (VP5) is surface exposed. We investigated whether the VP5 apical region contributes to intracellular transport dynamics following entry into primary sensory neurons and also tested the hypothesis that conserved negatively-charged amino acids in the apical region contribute to VP26 acquisition. To our surprise neither hypothesis proved true. Instead, mutation of glutamic acid residues in the apical region delayed viral propagation and induced focal capsid accumulations in nuclei. Examination of capsid morphogenesis based on epitope unmasking, capsid composition, and ultrastructural analysis indicated that these clusters consisted of procapsids. The results demonstrate that, in addition to established events that occur inside the capsid, the exterior capsid shell promotes capsid morphogenesis and maturation.

## IMPORTANCE

Herpesviruses assemble capsids and encapsidate their genomes by a process that is unlike other mammalian viruses but is similar to some bacteriophage. Many important aspects of herpesvirus morphogenesis remain enigmatic, including how the capsid shell matures into a stable angularized configuration. Capsid maturation is triggered by activation of a protease that cleaves an internal protein scaffold. We report on the fortuitous discovery that a region of the major capsid protein that is exposed on the outer surface of the capsid also contributes to capsid maturation, demonstrating that the morphogenesis of the capsid shell from its procapsid precursor to the mature angularized form is dependent upon internal and external components of the megastructure.

## INTRODUCTION

The *Herpesviridae* family of large double-stranded DNA viruses is divided into three subfamilies: the *Alpha, Beta*, and *Gammaherpesvirinae*. Several members of the *Alphaherpesvirinae*, including herpes simplex virus type 1 (HSV-1), exhibit robust neuroinvasive ability (1). HSV-1 infection begins in epithelial cells but quickly spreads to nerve terminals and invades the sensory and autonomic ganglia by retrograde axonal transport (2). Within these neuronal cells, HSV-1 establishes the lifelong, recurrent infection that is characteristic of these viruses. All herpesviruses share a common virion morphology consisting of three layers: nucleocapsid, tegument, and envelope. Herpesvirus nucleocapsids are approximately 125 nanometers in diameter, conform to a T = 16 icosahedral structure, and consist of 162 capsomeres: 150 hexamers, 11 pentamers, and one portal vertex (3). HSV-1 nucleocapsids are estimated to withstand 18 atmospheres of internal pressure exerted by the tightly packed 152 kilobase pair genome until the capsid docks at a nuclear pore and releases the DNA into the nucleus (4, 5). Delivery of the viral genome initiates a cascade of gene expression leading to the assembly of nascent virions.

Extensive efforts have been undertaken to understand herpesvirus capsid assembly and maturation, which includes analyses of baculovirus and cell-free systems (6–11), temperature-sensitive mutants (12, 13), and three-dimensional reconstructions from cryo-electron microscopy (14–20). From this work, an overarching model of HSV-1 capsid morphogenesis has been developed. The major capsid protein (VP5), triplexes (VP19c and VP23), portal vertex (pUL6), and internal scaffold proteins self-assemble into a spherical procapsid within the cell nucleus. There are two types of scaffold proteins: small (pUL26.5) and large (pUL26). Once assembly of the procapsid is complete the catalytic activity of the VP24 protease, housed within the amino terminus of the large scaffold protein, is triggered (21). VP24 cleaves itself from the scaffold and cleaves a release site in the C-terminus of the small and large scaffold proteins, disconnecting them from the shell floor (9, 22, 23). These events cause the capsid shell to angularize into a stable form and become a substrate for the binding of two accessory capsid surface proteins: pUL25 and VP26 (22, 24–26). The scaffold fragments internal to the capsid are expelled as the viral genome is packaged, but VP24 is retained (26–32). This process produces three angularized capsid types within infected cell nuclei: capsids that retained the scaffold and lack DNA (B capsids), capsids that expelled the scaffold but failed to stably encapsidate the viral genome (A capsids), and nucleocapsids (C capsids).

The majority of the capsid shell mass is comprised of 955 copies of the 149 kilodalton (kDa) VP5 protein (33–36), which is arranged into hexamers and pentamers. Pentamers are found at 11 of the capsid vertices while hexamers occupy the remainder of the icosahedral lattice, with the exception of the unique portal vertex. Hexamers are crowned with VP26 (12 kDa), and pentamers are associated with complexes of pUL25 and pUL17 that radiate outward from the pentamers and serve as tegument binding sites (5, 19, 37–40). The VP5 monomer is a large, tower-like protein divided into three domains: the upper, middle, and floor. The apical region of the upper domain, which consists of a polyproline loop plus an adjacent sequence that serves as an antibody binding epitope (41), is surface exposed in hexamers and pentamers and does not participate in subunit interactions within the capsid shell. Instead, the apical region is proposed to be important for binding VP26 on hexamers and tegument on pentamers (42).

Our studies of herpesvirus neuroinvasion demonstrate that capsid-bound tegument proteins contribute to neuronal transmission and retrograde axonal transport (43–45). To examine if the capsid surface participates in these processes, a collection of HSV-1 encoding mutations in the VP5 apical region was made. Initial characterization of these viruses in cell culture revealed only modest changes in propagation, with the exception of a virus mutated at two conserved acidic residues in the polyproline loop. Neither the polyproline loop mutant nor a mutant lacking the majority of the antibody epitope was impacted for retrograde axonal transport. Further examination of the polyproline loop mutant indicated that VP5 expression and VP26 incorporation onto capsids were also unimpaired. Using a combination of fluorescence and ultrastructural imaging, we determined that the polyproline loop mutant offers a unique example of a procapsid maturation defect that maps to the external face of the capsid shell.

## RESULTS

**Mutating VP5 apical region residues impacts herpes simplex virus spread in cell culture.** The majority of the HSV-1 major capsid protein, VP5, is integral to the capsid icosahedral structure and its constituent capsomeres. An exception is a small apical region within the VP5 upper domain that is exposed on the capsid surface (Fig. 1A-C) and consists of a polyproline loop and a binding site for a capsid-specific antibody (41, 42). The accessibility of the apical domain indicates it may interface with cellular or viral proteins to promote infection. To assess the importance of the apical region during HSV-1 infection, an alignment of eight alphaherpesviruses was used to inform a mutational approach (Fig. 1D). Based on sequence conservation, five VP5 mutants were made by two-step RED-mediated recombination in the F strain of HSV-1 (Fig. 1E). The recombinant viruses exhibited a reduction in cell-cell spread as evidenced by decreased plaque diameters compared to wild-type (WT) virus (Fig. 2A). The most severe decrease in plaque size occurred when two glutamic acid residues in the polyproline loop, E846 and E851, were mutated to alanines (EE>AA). These glutamic acid residues, shown on the VP5 crystal structure in magenta and blue in Figure 1A-D, are located in the most apical region of the VP5 monomer (Fig. 1C-D). The decrease in plaque size was only observed when both glutamic acid residues were mutated to alanine, and the defect was rescued when both mutations were repaired (Fig. 2B).

**Figure 1.**
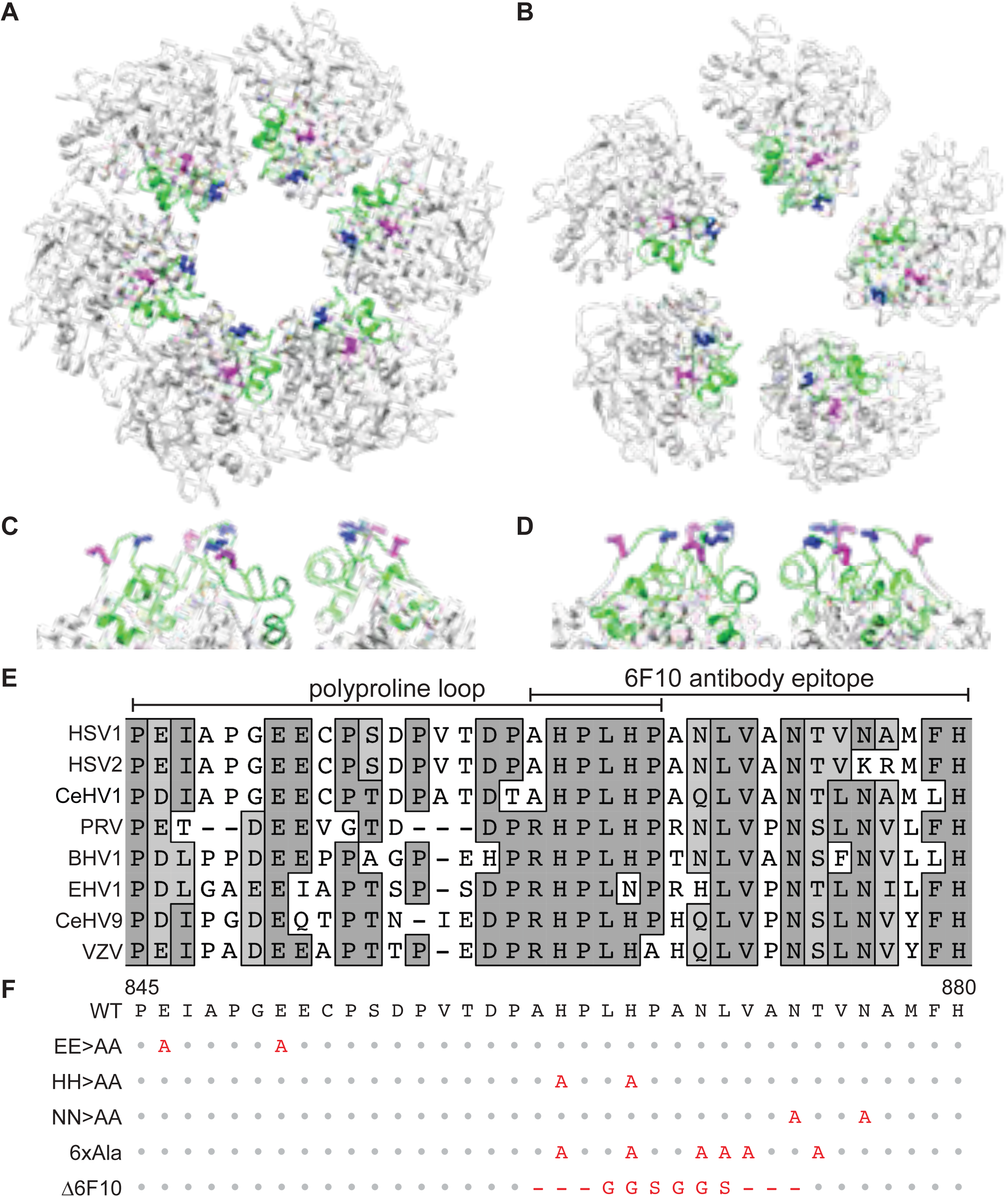
Targeted mutation of the VP5 apical region. (A-D) Representation of the HSV-1 VP5 upper domain crystal structures as arranged in a top view of (A) a peripentonal hexamer, (B) a pentamer, and a side view of (C) a peripentonal hexamer, and (D) a pentamer. The apical region is colored green and side chains of residues E846 (magenta) and E851 (blue) are shown as sticks. (E) Alignment of the apical region of eight alphaherpesvirus major capsid proteins. Amino acid positions for the HSV-1 sequence are indicated below. (F) HSV-1 VP5 apical region mutants used in this study. Predicted amino acid changes are in red.

**Figure 2.**
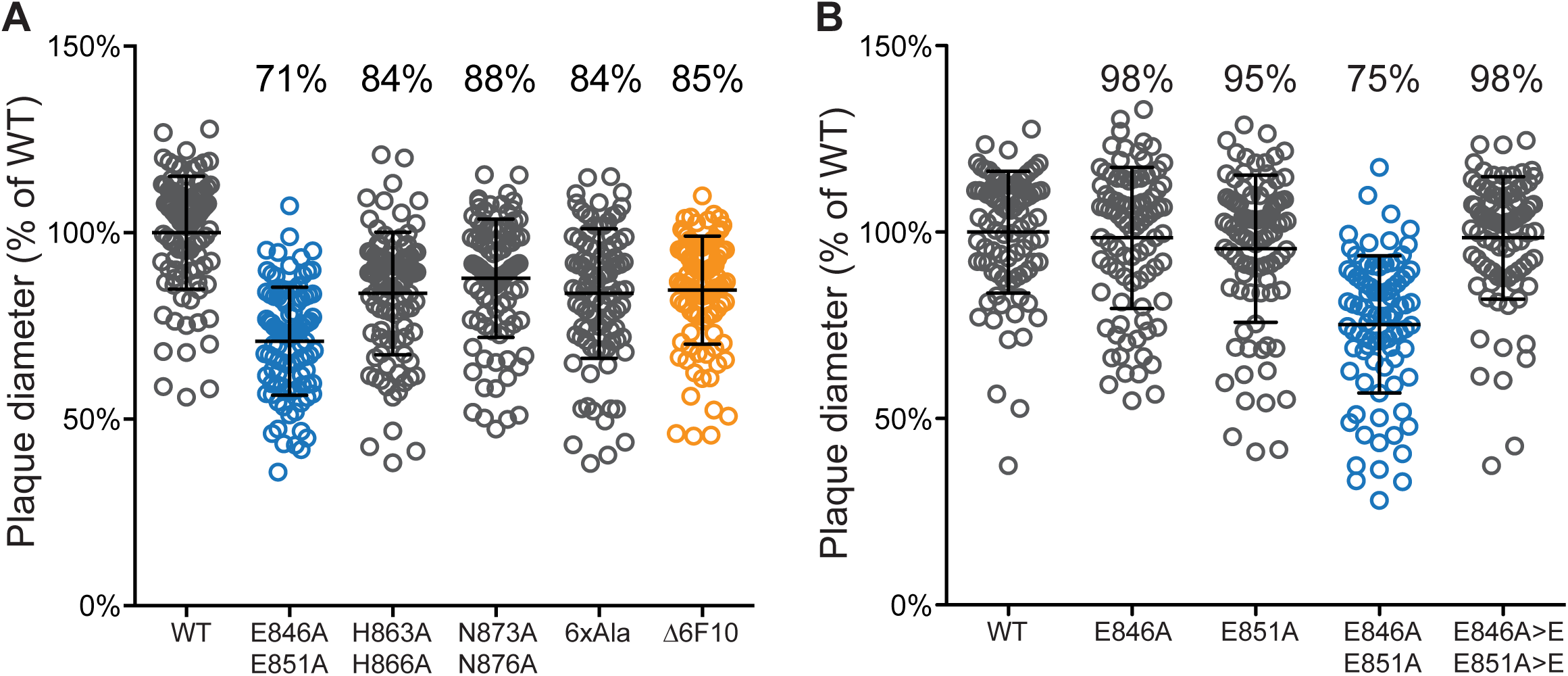
Impact of the VP5 mutations on plaque size. Vero cells were infected with (A) VP5 apical region mutants and (B) additional VP5 glutamic acid mutants. Plaque diameters were measured 5 days post infection. Diameters are represented as a percent of the mean plaque diameter of wild-type HSV-1 strain F (WT). Repaired alleles are listed as E846A>E and E851A>E. Error bars indicate the standard deviation of three independent experiments.

**The VP5 apical region does not contribute to retrograde viral transport.** The capsid-bound tegument proteins, pUL36 and pUL37, are effectors of intracellular capsid transport (5, 43–47). To examine if the capsid surface-exposed region of VP5 contributes to its transport on microtubules, primary sensory neurons were infected with the EE>AA and Δ6F10 mutant viruses (Fig. 1F). To visualize individual capsids in the process of retrograde axonal transport by time-lapse fluorescence microscopy, the viruses were further modified to encode a mCherry fusion to the maturation protease, VP24 (32). Capsid dynamics were monitored during the first hour post infection (hpi) and continuous periods of retrograde motion (runs) were analyzed for average velocity and distance traveled. The distribution of run velocities were consistently Gaussian for the wild-type, EE>AA, and Δ6F10 viruses (R^2^≥0.96 for each) and run distances were decaying exponential (R^2^≥0.98 for each), the latter being consistent with processive motion (Fig. 3). These transport dynamics were consistent between the WT and mutant viruses, demonstrating that mutation of the VP5 apical region did not impair microtubule-based retrograde axonal transport.

**Figure 3.**
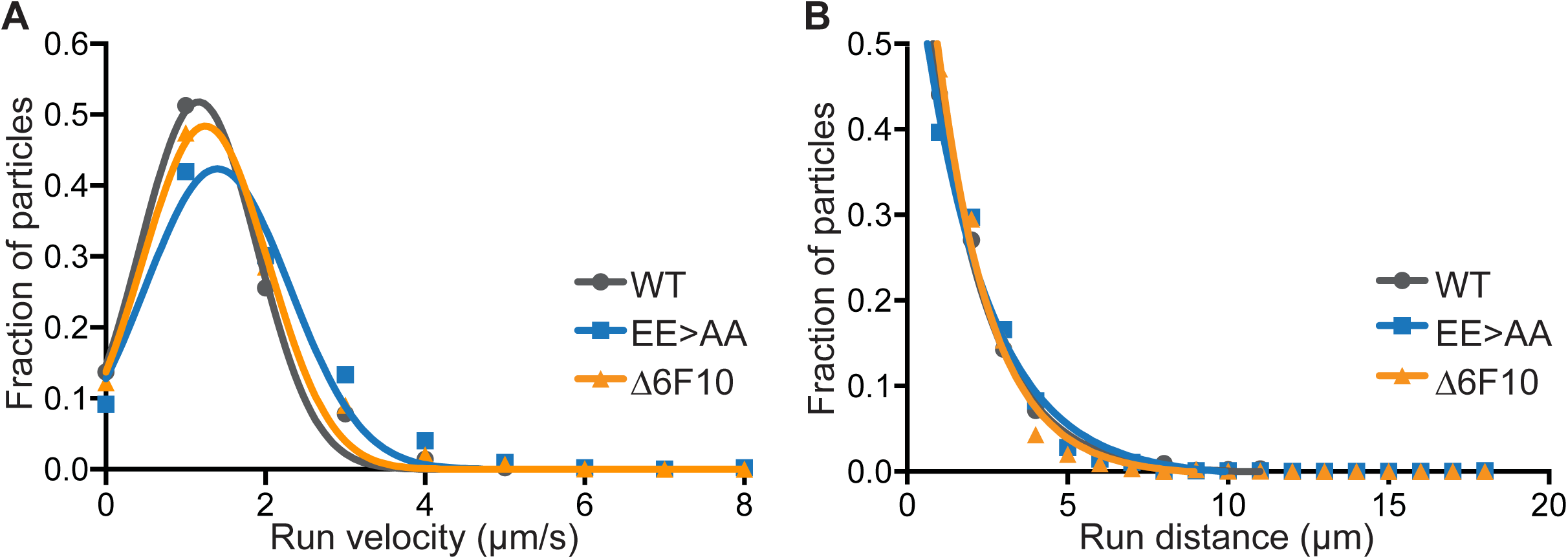
Mutation of the VP5 upper domain did not impair retrograde axonal transport. Primary sensory neurons were infected with derivatives of the VP5 wild-type (WT), glutamic acid mutant (EE>AA), and Δ6F10 mutant that encode a mCherry-VP24 fusion to allow for tracking of individual capsids in axons. Intracellular capsid transport was monitored by time-lapse fluorescence microscopy during the first hour of infection. Run velocity (A) and run distance (B) profiles of individual capsids are representative of three independent experiments.

**The VP5 EE>AA mutant forms intranuclear clusters of capsid proteins.** Because the defect underlying the reduction in plaque diameter was not ascribed to defects in retrograde trafficking, *de novo* capsid production was next examined. Vero cells infected with the mCherry-VP24 variants of the EE>AA and Δ6F10 mutants were examined by live-cell microscopy at 8.5 hpi. As expected, the nuclei of cells infected with either mutant virus filled with diffraction-limited punctae consistent with high loads of dispersed capsids (32); however, the EE>AA mutant also produced large intranuclear structures (Fig. 4A). These structures were also observed in the absence of the mCherry-VP24 tag but were not evident with WT HSV-1 or a repair of the EE>AA mutant (Fig. 4B), demonstrating that the EE>AA mutations induced intranuclear clusters of the capsid VP24 and VP5 capsid proteins. Because these foci were evident by a fusion protein and antibody staining, it suggests these foci were not artifacts of either the VP24 fluorescent tag or fixation.

**Figure 4.**
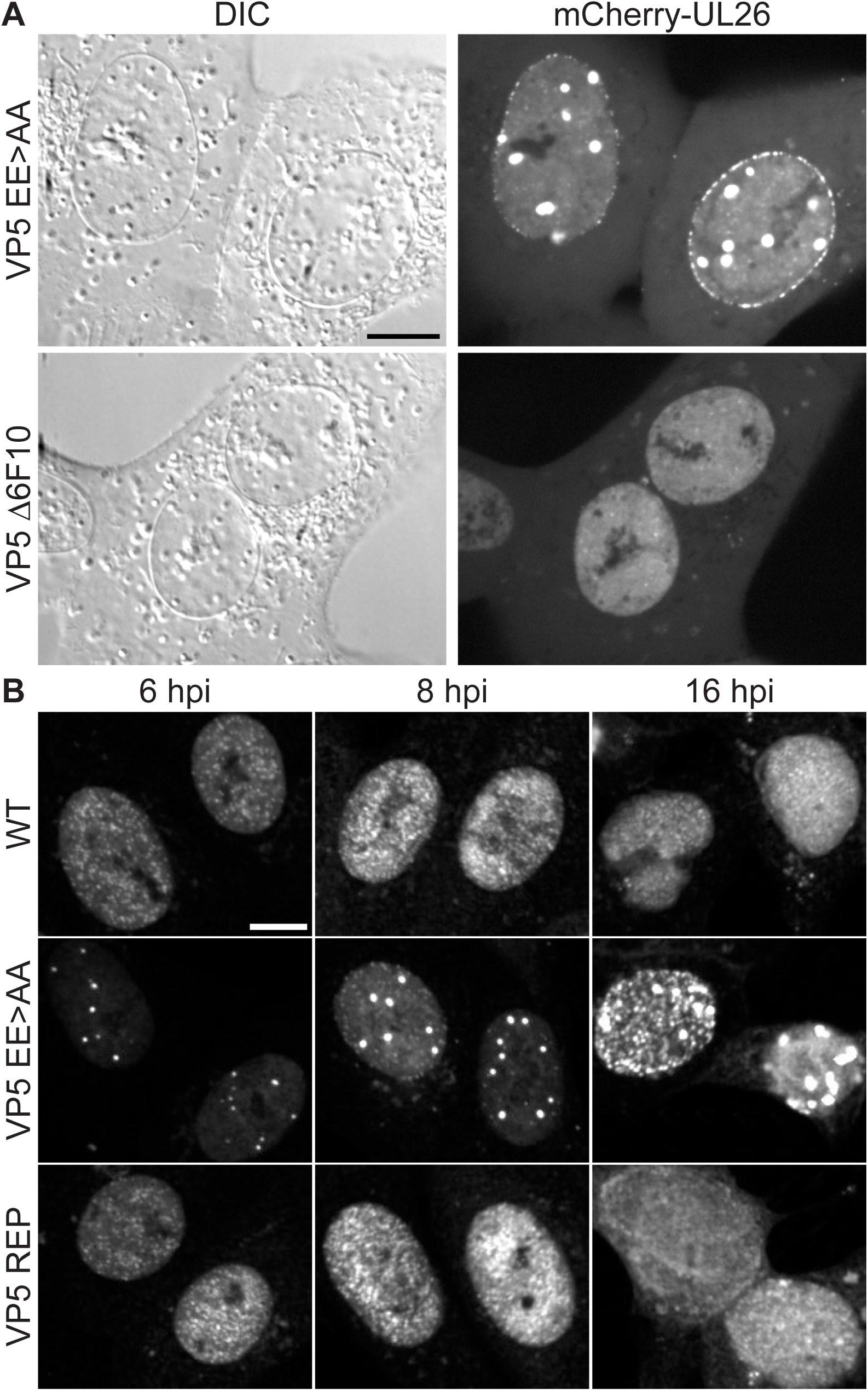
The VP5 EE>AA mutant forms intranuclear clusters of capsid proteins. Vero cells were infected at a MOI of 5. (A) Live-cell imaging at 8.5 hpi using derivatives of HSV-1 encoding mCherry-VP24 and either VP5 EE>AA or Δ6F10. (B) Fixed-cell immunofluorescence time course of cells infected with unmodified HSV-1 (WT), EE>AA mutant, and VP5 repair (REP). Cells were fixed at the indicated time post infection and stained with an aVP5 antibody. Scale bars are 10 μm.

**HSV-1 propagation kinetics are reduced by VP5 EE>AA mutation.** Single-step growth kinetics of the EE>AA mutant were attenuated relative to the wild-type and repair viruses (Fig. 5). At 8 and 12 hpi, the EE>AA mutant exhibited a decrease in both cell-associated and supernatant plaque forming units (PFU). By 24 hpi, the EE>AA mutant attained titers approaching the wild-type. We conclude that the aberrant nuclear foci correlated to decreased rates of propagation, as well as plaque size (Fig. 2), but had limited impact on the number of plaque-forming units per cell (burst size).

**Figure 5.**
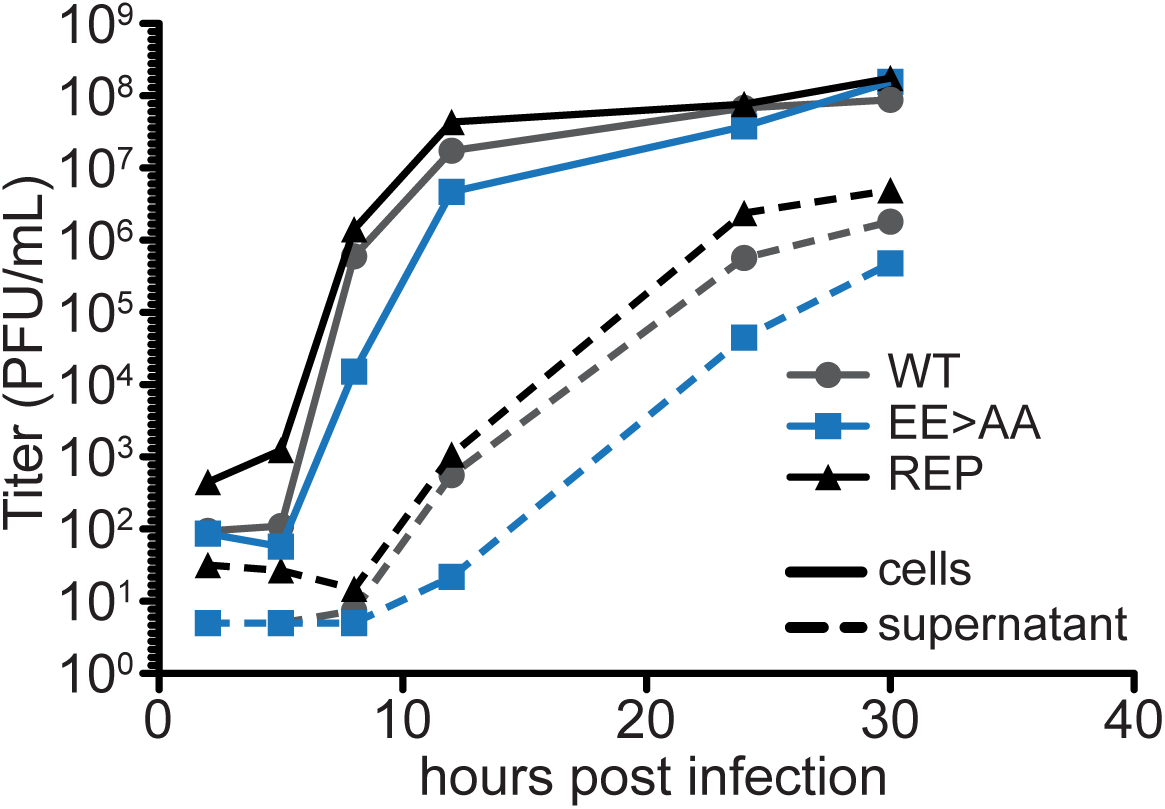
VP5 glutamic acid residues support HSV-1 propagation. Single-step replication kinetics of HSV-1 WT, VP5 glutamic acid mutant (EE>AA), and the VP5 repair in Vero cells are shown following infection at a MOI of 10. Cells (solid lines) and supernatants (dashed lines) were collected at indicated times and titers were determined by plaque assay on Vero cells.

**The conserved glutamic acids of the VP5 polyproline loop are dispensable for VP26 assembly.** Negatively-charged residues within the VP5 polyproline loop were predicted to mediate binding of the VP26 capsid protein onto the surface of capsid hexamers (42). To test this hypothesis, intranuclear capsids were isolated from Vero cells infected with wild-type HSV-1 (WT) or the glutamic acid mutant (EE>AA). A VP26-null mutant (ΔUL35) was included as a control. A, B, and C capsids were separated by density ultracentrifugation and visualized by light scattering. The EE>AA and VP26-null viruses produced each capsid species similar to the wild-type (Fig. 6A). The protein compositions of the A, B, and C capsids produced from the wild-type and EE>AA viruses were indistinguishable from one another; notably, capsids of the EE>AA virus possessed wild-type levels of VP26, which was identified by its 12 kDa molecular weight and its absence from the ΔUL35 samples (Fig. 6B). This data indicates that the VP5 EE>AA mutant assembles A, B, and C capsids and that E846 and E851 are dispensable for VP26 incorporation.

**Figure 6.**
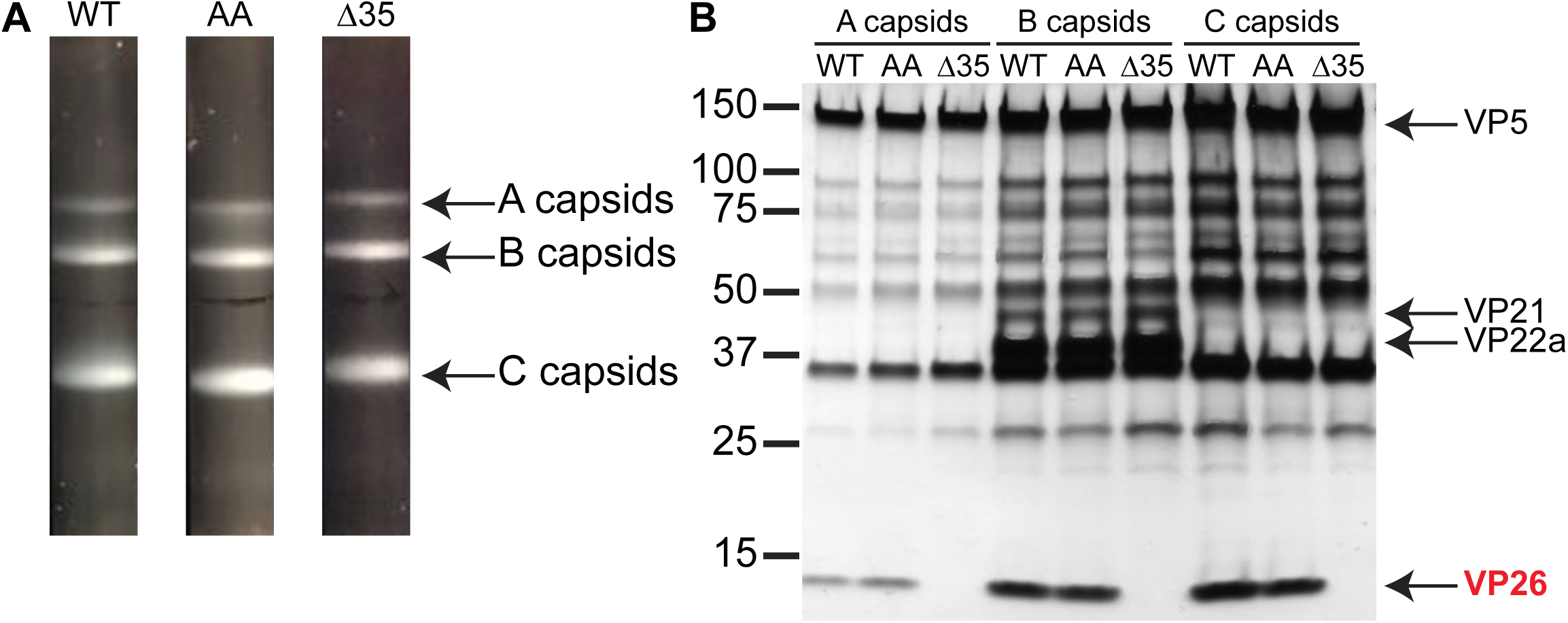
VP5 EE>AA capsids acquire VP26. A) A, B, and C capsids were isolated from nuclei of Vero cells infected with HSV-1 over a 20-50% sucrose gradient and visualized by light scattering. B) Protein profiles of the capsids observed by silver staining of a 4-20% gradient denaturing gel. HSV-1 strains used were wild-type (WT), the glutamic acid mutant (AA), and a ΔUL35 mutant (Δ35). The latter serves to identify VP26 in the denaturing gel.

**The VP5 EE>AA mutant produces wild-type levels of VP5 protein**. The capsid profiles from the previous experiment were consistent with the expression of capsid proteins and their assembly. Nevertheless, the potential impact of the EE>AA mutation on VP5 expression warranted further inspection. Western blot detection of VP5 from whole-cell lysates was performed at 8 hpi. VP5 was detected as a 149 kDa band in both HSV-1 WT and EE>AA but not mock-infected cell lysates (Fig. 7A). The expression of VP5 was normalized to the pUL37 tegument protein as a loading control following densitometry analysis, with wild-type HSV-1 set to a value of 1. The analysis revealed no significant change in the amount of VP5 produced by the EE>AA mutant (Fig. 7B). Therefore, we next examined the impact of EE>AA mutation on capsid morphogenesis.

**Figure 7.**
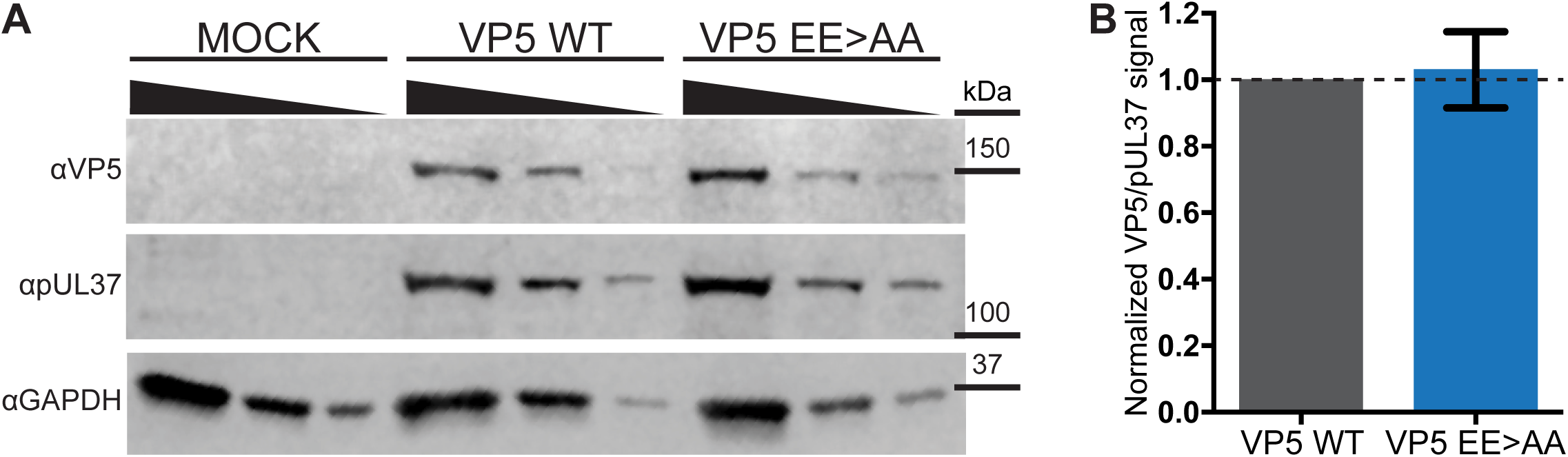
VP5 expression is not impacted by glutamic acid mutations. (A) Western blots of cell lysates prepared from mock or infected cells. Vero cells were infected with HSV-1 encoding the wild-type (WT) or EE>AA VP5 allele at a MOI of 5. Threefold dilutions of cell lysates, prepared at 8 hpi, were probed by western blot. The HSV-1 tegument protein, pUL37, and cellular GAPDH served as loading controls. The image shown is representative of three independent experiments. (B) VP5 expression was quantitated by densitometry analysis. Individual measurements are normalized to the pUL37 loading control, and the average expression levels are presented relative to the WT sample. The difference was not statistically significant by an unpaired t-test. Error bars represent the standard error of the mean.

**Nuclear foci formed during infection with the HSV-1 VP5 EE>AA mutant initially lack capsid maturation markers.** To better understand the protein composition of EE>AA foci, Vero cells were infected, fixed at 8 hpi, and processed for fluorescence imaging. Cells infected with HSV-1 encoding wild-type VP5 had capsids dispersed throughout the nuclei, which substantially colocalized with VP24, VP26, pUL25, and with reactivity to the maturation-specific VP5 antibody, 8F5 (Fig. 8). In contrast, the EE>AA nuclear foci colocalized only with VP24. Because the VP26 and pUL25 capsid proteins are acquired upon shell angularization (22, 24–26), and the VP5 8F5 epitope (48) is exposed after internal scaffold cleavage by the maturation protease (49, 50), we conclude the EE>AA nuclear foci did not contain mature capsids. However, by 16 hpi, the foci acquired each of these markers (Fig. 9).

**Figure 8.**
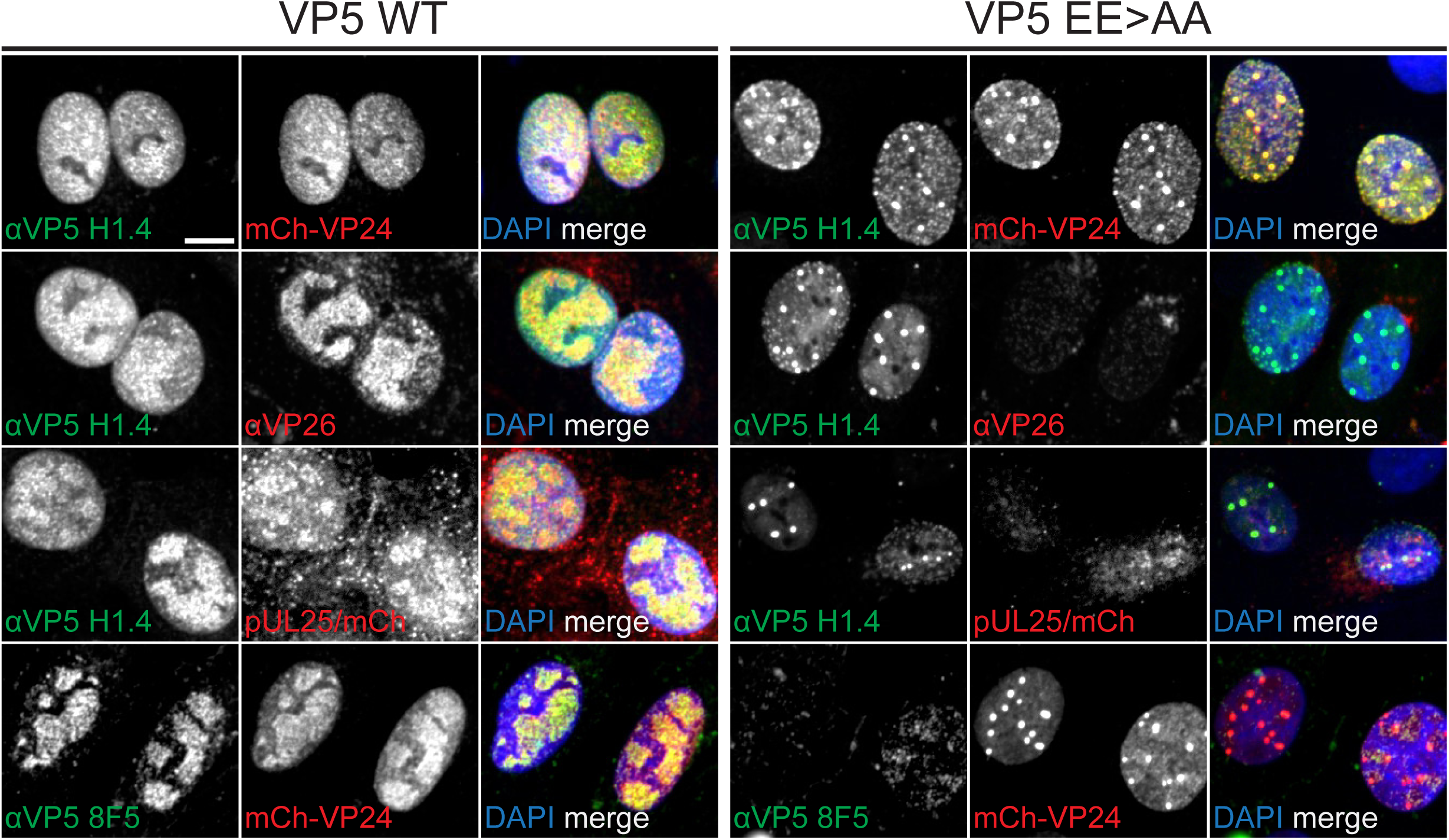
VP5 EE>AA foci composition is consistent with that of procapsids. Vero cells infected with HSV-1 encoding WT or EE>AA VP5 at a MOI of 5 were fixed at 8 hpi and processed for immunofluorescence using aVP5 and αVP26 antibodies. In some instances, the viruses encoded mCherry-VP24 or pUL25/mCherry fusions, as indicated. Scale bar is 10 μm.

**Figure 9.**
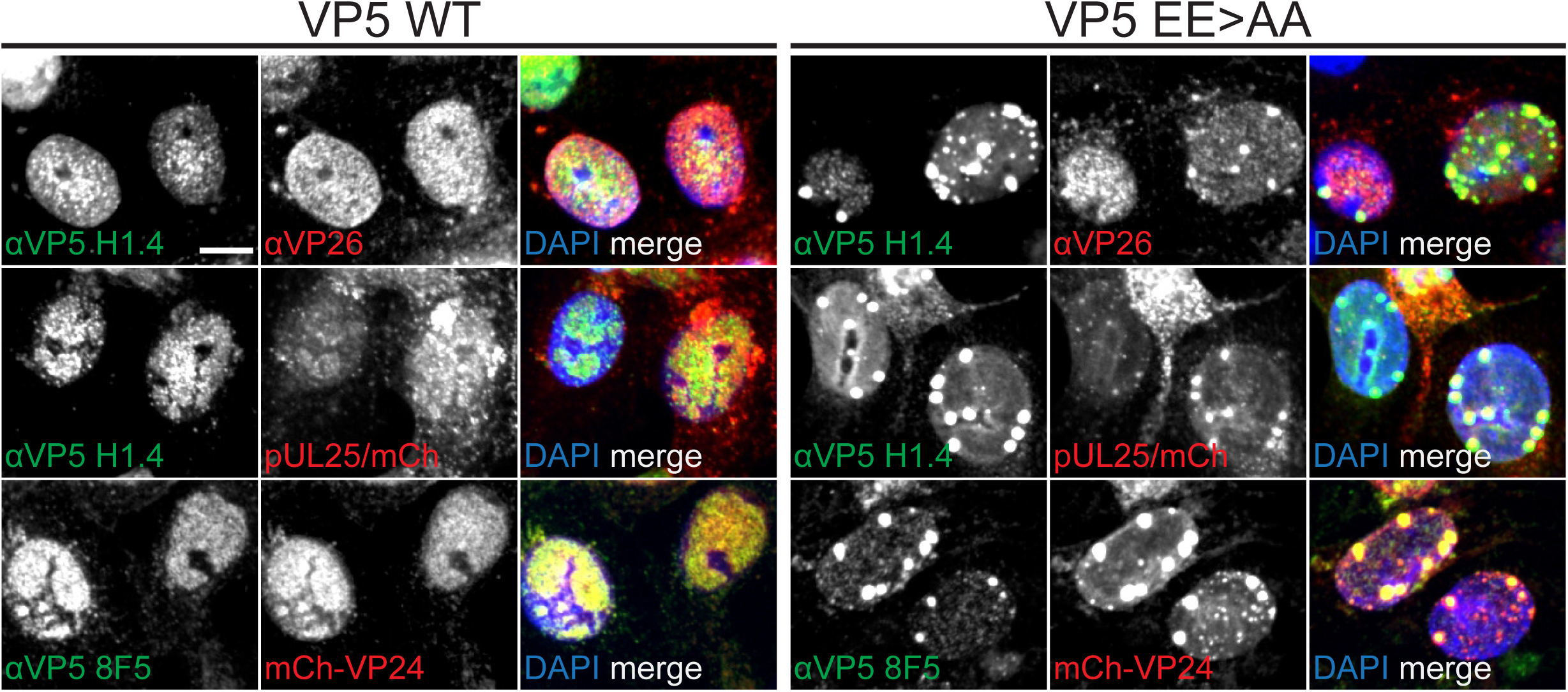
VP5 EE>AA foci acquire capsid maturation markers during late infection. Experiments were carried out as described for Figure 7 but were examined at 16 hpi. Scale bar is 10 μm.

**The VP5 EE>AA foci contain procapsids.** Transmission electron microscopy was used to visualize the EE>AA foci. Vero cells infected with HSV-1 VP5 EE>AA and fixed at 8 hpi contained nuclear foci of electron-dense material surrounding partially- and fully-formed procapsids (Fig. 10A and B). Consistent with the production of infectious EE>AA virions (Fig. 5), mature capsid species were detected throughout these nuclei but always apart from the foci (Fig. 10C-F). At 16 hpi, the intranuclear foci persisted but now contained mixtures of procapsids and B capsids (Fig. 10G-H). Interestingly, only scaffold-containing capsids were observed within the VP5 EE>AA foci with C capsids found outside the foci (Fig. 10I). Collectively, the data indicate that mutation of the E846 and E851 residues on the capsid surface delays procapsid maturation.

**Figure 10.**
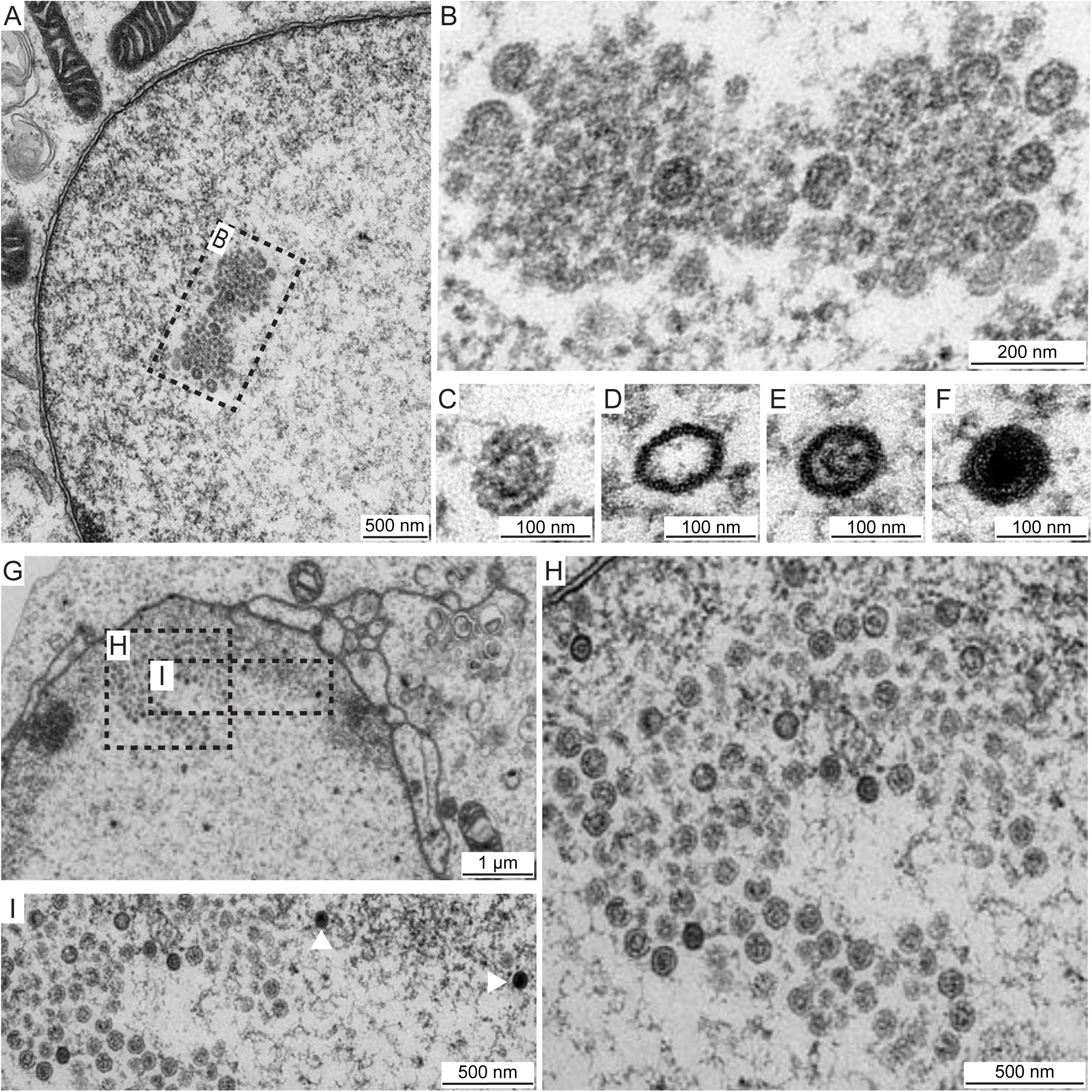
VP5 EE>AA foci contain clusters of procapsids. HSV-1 VP5 EE>AA infected Vero cells (MOI of 5) were fixed at 8 hpi (A-F) or 16 hpi (G-I) and imaged by transmission electron microscopy. A and B) Foci ultrastructure reveals fully- and partially-formed capsids containing large-core scaffolds consistent with procapsids. C, D, E, F) Infected cells exhibit all capsid species: C) procapsid, D) A capsid, E) B capsid, and F) C capsid. Scale bars are 100 nm (C-F). G and H) Procapsids persist in foci and B capsids are now evident as small-core capsids. I) C capsids (white arrowheads) are present in nuclei but are excluded from foci.

## DISCUSSION

Assembly of herpesvirus virions begins with the formation of spherical procapsid shells in cell nuclei. Procapsids are short-lived structures that are unstable when isolated from infected cells, but spontaneously angularize into rigid, stable capsids during maturation (13, 32). When shell maturation is accompanied by genome encapsidation, the resulting product is a C capsid, or nucleocapsid. This critical process is a target for antiviral development (51–59), which is bolstered by ongoing refinements of cryo-EM reconstruction techniques that have led to herpesvirus capsid models with near-atomic resolution (19, 60–62).

The surface of the procapsid is decorated by the apical region of the VP5 major capsid protein, which constitutes the tips of the 11 pentamers (accounting for 11 of the icosahedral vertices, with the 12^th^ vertex consisting of 12 copies of pUL6 portal protein) and 150 hexamers that make up the majority of the icosahedral shell. Following maturation, the procapsid acquires the pUL25 and VP26 accessory proteins, the latter of which selectively covers the apical regions of the hexamers. Thus, the procapsid displays 955 copies of the apical region on its surface, which is reduced to 55 exposed copies upon maturation and decoration with VP26. The exposure of the VP5 apical region on the mature capsid led us to investigate whether it contributes to the retrograde axonal transport of capsids that leads to the establishment of latent infections in the nervous system.

Although no functions have formally been ascribed to the apical region, two conserved glutamic acids within the polyproline loop of the apical region (E846 and E851) were proposed as potential sites for recruitment of VP26 upon shell maturation (18, 42). While residues within the VP5 upper domain are undoubtedly important for the VP26-VP5 interaction, our data demonstrated that the two glutamic acid residues are not required for VP26 binding to capsids. Furthermore, they were dispensable for retrograde axonal transport in primary sensory neurons, as was the antibody epitope portion of the apical region (based on analysis of the Δ6F10 mutant). Given that the VP26 protein is also dispensable for axonal transport (63, 64), we conclude that the bulk of exposed outer capsid surface does not promote this critical component of the HSV-1 neuroinvasion mechanism. While the scope of the mutagenesis included in this study does not entirely rule out a role for capsid surface epitopes that contribute to the transport mechanism, it seems likely that the previously identified interaction between pUL25 and the pUL36 tegument protein may be sufficient to confer these dynamic properties to the virus particle (65). These results support a model in which the herpesvirus capsid is a dormant cargo that requires the action of attached tegument proteins to produce the dynamics that underlie productive infection.

Despite these findings, the polyproline glutamic acid mutant produced small plaques in cell culture, indicating the apical region was not fully dispensable. Further analysis correlated the propagation defect with the formation of focal structures in infected cell nuclei that consisted of clusters of procapsids. This observation was unexpected because it indicated that capsid maturation was delayed: a process dependent on proteolytic processing of the interior scaffold. While the VP5 floor domain faces the interior of the capsid and interacts with the scaffold proteins (66–70), the exterior location of the apical region is inconsistent with it promoting scaffold processing. Furthermore, the VP5 apical region does not participate in inter-capsomeric interactions (14, 15, 17–19, 62). Importantly, the persistence of procapsids that was initially indicated by the absence of maturation markers (pUL25, VP26, and exposure the 8F5 epitope) was verified by ultrastructural analysis. The latter revealed the foci to consist of capsids containing large scaffold cores, which are indicative of procapsids (7, 13, 71). While the acquisition of surface markers could be sterically restricted within foci, the large scaffold cores indicate that the VP24 maturation protease was inefficiently activated within these procapsids: an event that does not require any known extrinsic interactions including with pUL25, VP26 or genome encapsidation (7, 32, 72, 73).

We note that the procapsid foci of the polyproline mutant were associated with electron-dense material when imaged by transmission electron microscopy, and in this way resembled procapsids in cell-free assembly systems (9, 74). The procapsid foci were also reminiscent of foci observed when VP24 activity is disrupted through mutation or deletion (12, 13, 75). In contrast, we previously reported a catalytically-inactive VP24 mutant that did not produce procapsid foci (32). Thus, the mechanism driving procapsids to cluster in foci is of interest, as the clusters may represent sites of capsid assembly. Of greater interest is why capsid maturation is sensitive to mutations in the polyproline loop of the VP5 apical region. While procapsids in the mutant foci were often seen in a partially assembled state, the defect is inconsistent solely with an assembly defect, as seeing accumulations of procapsids is atypical of wild-type infections. Because angularization of the capsid shell under physiological conditions is dependent solely on VP24 activation, which occurs independently of genome encapsidation (9, 32, 76, 77), we can infer that mutation of the apical region has affected VP24 activation. Although speculative, the results are consistent with VP24 activation being enhanced by a cue external of the capsid. Capsids require ATP for shell maturation (78), and the apical domain could respond to ATP or interface with another factor such as the putative chaperone of preassembled capsids, pUL32 (79). Interestingly, the mutant foci exclusively consisted of procapsids and B capsids, with procapsids being the predominant species at 8 hpi and B capsids accumulating by 16 hpi. While A and C capsids could be found in proximity to the foci, there was no evidence of DNA-packaging occurring within them.

Despite a wealth of knowledge, the details of the molecular mechanisms driving procapsid assembly and maturation remain enigmatic. Viral mutants are a valuable resource to better understanding the complex assembly and maturation of nucleocapsids (13, 78). To the best of our knowledge, this report presents the first attribution of function to the VP5 apical region and documents that the exterior of the capsid contributes to shell maturation.

## MATERIALS AND METHODS

**Sequences and alignment.** Major capsid protein amino acid sequences obtained from GenBank are listed in order of appearance from top to bottom of the alignment in Figure 1E: GU734771, Z86099, NC_004812, JF797219, AJ004801, AY665713, NC_002686.2, and NC_001348.1. Protein sequences were aligned using the ClustalW alignment tool in MacVector.

**Structural modeling.** Using UCSF chimera (80) the structure of the upper domain of VP5 (PDB: 1NO7) (42) was fit in both the pentamer and the peripentonal hexamer of the previously published density map of the HSV-1 (EMDB: EMD-6386) (18).

**Cells and virus.** Vero (African green monkey kidney epithelial) and Vero cells expressing Cre recombinase (Vero-CRE) were maintained in Dulbecco’s Modified Eagle Medium (DMEM) (Invitrogen, Catalog #11965-118) supplemented with 10% bovine growth serum (BGS) (vol/vol) (Rocky Mountain Biologicals, Catalog #FGR-BBT). The latter were generously provided by Dr. David Leib (81). Cultured cells were tested for *Mycoplasma* contamination with the PlasmoTest Mycoplasma Detection Kit (InvivoGen, Catalog #rep-pt1). Primary sensory neurons were isolated from the dorsal root ganglion (DRG) of embryonic chicks (E9-10) and cultured as whole explants as previously described (82). Table 1 details all recombinant viruses used in this study. Mutation of UL19, UL35, or insertion of coding sequences for fluorescent proteins was achieved by a two-step RED-mediated recombination method (83). Primers used for BAC mutagenesis are listed in Table 2. Mutations were sequence confirmed at the Northwestern University Genomics Core Facility. HSV-1 recombinant viruses were produced by electroporation of the infectious bacterial artificial chromosome (BAC) clone of HSV-1 strain F, pYEbac102, into Vero cells (84). Recombinant viruses were subsequently propagated on Vero-CRE cells to excise the loxP-flanked BAC backbone from the viral genome and produce a working stock of virus. All viral stock titers were determined by plaque assay on Vero cells overlaid with DMEM supplemented with 10 mg/ml methyl cellulose (VWR, Catalog #AA36718-36), Penicillin-Streptomycin (Pen-Strep) at 100 U/mL (Invitrogen, Catalog #15140122), and 2% BGS.

**TABLE 1.** Recombinant viruses

| Strain | Fluorescent fusion | Mutations | Titer (PFU/mL) |
| --- | --- | --- | --- |
| HSVF-GS2695<sup>a</sup> | - | - | $2.59 \times 10^8$ |
| HSVF-GS5798 | - | UL19 E846A/E851A | $4.72 \times 10^7$ |
| HSVF-GS5799 | - | UL19 N873A/N876A | $9.81 \times 10^7$ |
| HSVF-GS5801 | - | UL19 H863A/H866A | $1.31 \times 10^8$ |
| HSVF-GS5905 | - | UL19<br>H863A/H866A/N869A/L870A/V871A/T874A | $7.56 \times 10^7$ |
| HSVF-GS6276 | - | UL19 A862-N873 > GGSGGS | $1.07 \times 10^8$ |
| HSVF-GS6197 | - | UL19 E846A | $1.45 \times 10^8$ |
| HSVF-GS6434 | - | UL19 E851A | $1.93 \times 10^8$ |
| HSVF-GS6462 | - | UL19 E846A>E/E851A>E | $3.06 \times 10^8$ |
| HSVF-GS5748 | mCherry-pUL26 | - | $2.95 \times 10^8$ |
| HSVF-GS6460 | mCherry-pUL26 | UL19 E846A/E851A | $8.06 \times 10^7$ |
| HSVF-GS6493 | mCherry-pUL26 | UL19 A862-N873 > GGSGGS | $1.32 \times 10^7$ |
| HSVF-GS4553<sup>b</sup> | pUL25/mCherry | - | $7.65 \times 10^7$ |
| HSVF-GS5797 | pUL25/mCherry | UL19 E846A/E851A | $3.70 \times 10^7$ |
| HSVF-GS6199 | - | ΔUL35 | $1.59 \times 10^8$ |

<sup>a</sup> Previously published in (88)<sup>b</sup> Previously published in (45)

**TABLE 2.**
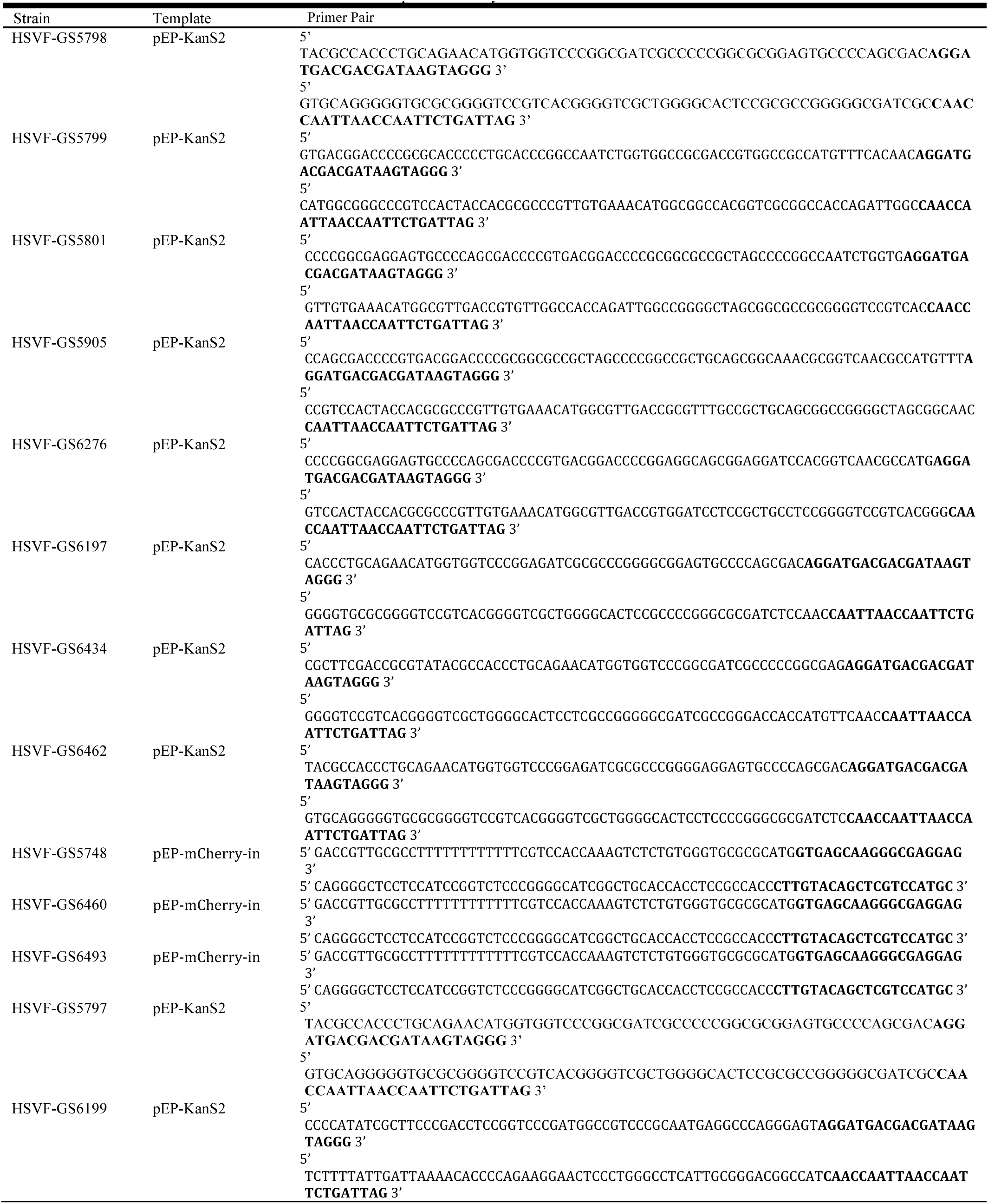
Primers used for RED-mediated recombination unique to this study

**Virus propagation and plaque assays.** Single-step growth curves were determined as previously described (85) with the following changes. In brief, Vero cells seeded in 6-well plates were infected at a multiplicity of infection (MOI) of 10. After 1 hour, unabsorbed virus was inactivated with 1 ml of citrate buffer (pH 3.0), cells were washed, and then incubated in 2 mL of DMEM supplemented with 2% BGS at 37°C, 5% CO_2_. Cell-associated and extracellular virus was harvested at 2, 5, 8, 12, 24, and 30 hpi. Titers were determined in duplicate by thawing and sonicating samples and performing plaque assay on Vero cells.

Plaque assays used for plaque diameter measurements were incubated 5 days post infection and then stained with Neutral Red (Santa Cruz, Catalog #sc-281691). Cells were gently rinsed with PBS before adding 1 mL of 1:1 Neutral Red and PBS. After 1 hour, stain was replaced with PBS and plaques were imaged on an Epson Perfection V500 Photo scanner at 3,200 dots per inch. Plaque diameters were determined by averaging two orthogonal diameter measurements for each individual plaque. Plaque diameters were then expressed as percent of wild-type (WT) plaque diameter and plotted in Prism 7 (Graphpad). More than 85 plaques were measured for each virus.

**Live-cell fluorescence microscopy.** To monitor capsid retrograde axonal transport, whole-explant chick DRGs were infected, imaged, and analyzed as previously described (86) with the following changes. Imaging began approximately 30 minutes after infecting DRGs with 3-5 × 10^7^ plaque forming units (PFU). Live-cell images were captured on an inverted wide-field Nikon Eclipse TE2000-U microscope fitted with a 60x/1.40 numerical aperture (NA) objective and a Cascade II:512 camera. Run-velocities and run-distances traveled were determined by kymograph analysis. Viral particles with a forward (retrograde) motion > 0.5 μm were plotted on a histogram as fraction of total particles and run-velocities or run-distances were fit to a Gaussian or decaying exponential curve with a non-linear regression, respectively. More than 150 runs were measured for each virus.

For live-cell imaging of infected epithelial cells, Vero cells were seeded on flame-sterilized 22 × 50 mm No. 1.5 cover glass and infected at a MOI of 5. After 1 hour, inoculum was replaced with F12 media supplemented with 2% BGS and Pen-Strep. VALAB chambers were made on 25 × 75 × 1mm slides as previously described (82) at 8.5 hpi – a time when viral capsids can readily be observed in infected cell nuclei (13, 32). Slides were imaged with a 100x lambda/1.49 NA objective on a Nikon Ti inverted microscope housed in an environmental box set at 37°C (InVivo Scientific) and coupled with a CSU-W1 confocal head (Yokogawa Electric Corporation) and a CascadeII:1024 EM-CCD (Photometrics). Illumination was supplied by a Sapphire 561 nm laser (Coherent) and custom laser launch (Solamere Technology Group, Inc.).

**Preparation of whole-cell lysates.** Vero cells seeded in a 10-centimeter (cm) dish were infected at a MOI of 5. After 1 hour, inoculum was replaced with DMEM supplemented with 2% BGS. At 8 hpi, cells were washed with 10 mL of ice cold PBS and lysed in 300 μL of RIPA buffer [50 millimolar (mM) Tris pH 8, 150 mM NaCl, 1% NP40, 0.5% sodium deoxycholate, 0.1% sodium dodecyl sulfate] supplemented with 5 mM dithiothreitol and protease inhibitors [2.5 mM sodium fluoride, 1 mM sodium orthovanadate, 0.5 mM phenylmethylsulfonyl fluoride, 100 μL of protease inhibitor cocktail (Sigma, Catalog #P8340)]. Cells were scraped into lysis buffer, transferred to an Eppendorf tube, and set on ice for 15 minutes. Lysates were then sonicated for three 1.5 second pulses and set on ice for an additional 15 minutes. Lysates were spun at 13,201 × g for 20 minutes and the supernatants were harvested. Total protein concentration of whole-cell lysates was determined using the Pierce 660 nm Protein Assay Reagent (Thermo Fisher Scientific, Catalog #22660). Absorbances of lysates was read at 650 nm wavelength on a Molecular Devices SpectraMax M5 Microplate Reader. Sample protein concentration was calculated from the standard curve of Pierce Pre-Diluted Protein Assay Standards: Bovine Serum Albumin (BSA) Set (Thermo Fisher Scientific, Catalog #23208). Samples were diluted in 5x final sample buffer (FSB) [60 mM Tris HCl pH 6.8, 2% SDS, 10% glycerol, 0.01% bromophenol blue] supplemented with 5% β-mercaptoethanol and stored at −20°C.

**Western blot and densitometry analysis.** Whole-cell lysates were thawed, boiled for 5 minutes, and loaded as three three-fold dilutions in equal concentrations across samples on a 4-20% mini-PROTEAN TGX Gel (Catalog #456-1096). Bio-Rad Dual Color standards (Bio-Rad, Catalog #1610374) were also loaded to mark molecular weights. Proteins were separated and transferred to a PVDF membrane (Millipore, Catalog #IPFL00010). The membrane was blocked in 5% milk for 80 minutes and then incubated overnight with primary antibodies diluted in 1% milk in PBST (0.1% Tween 20 in PBS) as follows: mouse monoclonal anti-VP5 H1.4 (Meridian Life Science, Inc., Catalog #C05014MA) at 1:1000 and rabbit polyclonal anti-pUL37 (kindly provided by Frank Jenkins) at 1:500. The next day, the membrane was washed and incubated for 1 hour at room temperature (RT) with secondary antibodies diluted in 1% milk in PBST as follows: goat anti-mouse IRDye 800CW (LI-COR Biosciences, Catalog #926-32210) at 1:10,000 and donkey anti-rabbit IRDye 680RD (LI-COR Biosciences, Catalog #926-68073) at 1:10,000. The membrane was imaged on a LI-COR Odyssey FC imaging system at 700 nm and 800 nm wavelengths for 3 minutes and 30 seconds each wavelength. Next, the membrane was blocked again with 5% milk in PBS for 80 minutes and then incubated for 1 hour at RT with primary antibody mouse monoclonal αGAPDH [clone 6C5] (Abcam, Catalog #ab8245) diluted 1:500 in 1% milk in PBS. The membrane was washed and then incubated at RT for 1 hour with secondary antibody goat amouse IRDye 800CW at 1:10,000 and imaged on the LI-COR Odyssey FC imaging system at 700 nm for 3 minutes and 30 seconds. Densitometry analysis was carried out using Image Studio Lite software. In brief, rectangles were added around each VP5 and pUL37 signal band with background subtraction set to median with a border width of 3 pixels on top and bottom. Boxes were adjusted, if needed, based on the profiles shape and intensity tools. The signal values for VP5 bands were calculated as a ratio to their corresponding pUL37 signal value. These ratio values were then normalized to wild-type samples.

**Immunofluorescence assays.** Vero cells seeded on flame-sterilized cover glass were infected at a MOI of 5 for 1 hour and then incubated in DMEM supplemented with 2% BGS at 37°C, 5% CO_2_. At the indicated time post infection, infected cells were washed with PBS and then fixed in 4% paraformaldehyde for 15 minutes. Fixed cells were rinsed three times with PBS and then permeabilized in PBST (0.1% Triton X-100 in PBS) for 10 minutes. Permeabilized cells were blocked in 5% bovine serum albumin (BSA) (Sigma, Catalog #A9647) in PBST for 45 minutes at RT or overnight at 4°C. After blocking, fixed cells were reacted with primary and secondary antibodies each for 1 hour at RT protected from light. Primary antibodies were diluted in 1% BSA in PBST as follows: mouse monoclonal anti-VP5 H1.4 (Meridian Life Science, Inc., Catalog #C05014MA) at 1:30, mouse monoclonal anti-VP5 8F5 (kindly provided by Duncan Wilson) at 1:100, and rabbit anti-VP26 (kindly provided by Prashant Desai) at 1:2000. Secondary antibodies were diluted in 1% BSA in PBST as follows: goat anti-mouse Alexa Fluor 488 (Thermo Fisher Scientific, Catalog #A-11001) at 1:400 and goat anti-rabbit Alexa Fluor 568 (Thermo Fisher Scientific, Catalog #A-11011) at 1:400. Cover glass was mounted on slides with ProLong Gold with DAPI mounting media (Thermo Fisher Scientific, Catalog #P36931). Slides were imaged by confocal microscopy on a Nikon Ti inverted microscope fitted with a 60x/1.49 NA objective as described above.

**Transmission electron microscopy.** Vero cells seeded in a 10-centimeter dish were infected at a MOI of 5 for 1 hour and then incubated in DMEM supplemented with 2% BGS and Pen-Strep at 37°C, 5% CO_2_. At the indicated time post infection, infected cells were rinsed with PBS and fixed in 2.5% glutaraldehyde in 0.1 M cacodylate buffer for 60 minutes at RT. After fixation, cells were scraped into fixative and transferred to an Eppendorf tube. Fixed cells were pelleted at 300 x g for 10 minutes. The cell pellet was enrobed in 5% low-melting point agarose (SeaPlaque^®^ Agarose, Catalog #50101) and delivered to the Northwestern University’s Center for Advanced Microscopy to be processed for transmission electron microscopy. In brief, the agarose plug was post-fixed with 1% Osmium tetroxide, stained with 0.5% uranyl acetate, dehydrated in ascending grades of ethanol from 30-100%, infiltrated with propylenoxide and embedded in hard resin. Blocks were thin sectioned at 50-60 nm and stained with uranyl acetate and Reynolds lead citrate. Slices were mounted on 200 mesh copper grids (Electron Microscopy Sciences). Imaging work was performed on a FEI Tecnai Spirit G2 transmission electron microscope at the Northwestern University Center for Advanced Microscopy generously supported by NCI CCSG P30 CA060553 awarded to the Robert H Lurie Comprehensive Cancer Center.

**Intranuclear capsid isolation and silver staining.** Isolation of intranuclear capsids was performed as previously described (87) with the following changes. Vero cells were seeded in 15-cm dishes and infected at a MOI of 10 with five dishes per virus. Infected cells were harvested between 22-24 hpi. A, B, and C capsid bands were pulled from the gradient using a Gradient Fractionator (BioComp Instruments). Each capsid species was subsequently diluted in a total volume of 2 mL of TNE [20 mM Tris pH 7.6, 500 mM NaCl, 1 mM EDTA] supplemented with 10 μL of protease inhibitor cocktail (Sigma, Catalog #P8340) and pelleted in a Beckman SW50.1 rotor at 74,909 x g for 1 hour and 4 minutes at 4°C. Pelleted capsids were resuspended in 50 μL of 5x FSB supplemented with 5% β-mercaptoethanol and stored at −20°C.

Resuspended capsids were thawed, boiled for 5 minutes, and loaded in equal volumes on a 4-20% Mini-PROTEAN TGX Gel (Catalog #456-1096) with Bio-Rad Dual Color standards (Bio-Rad, Catalog #1610374) diluted 1:100 in 5x FSB supplemented with 5% β-mercaptoethanol. Proteins were then detected using the Pierce Silver Stain Kit (Thermo Fisher Scientific, Catalog #24612). The silver stained gel was imaged on an Epson Perfection V500 Photo scanner at 2400 dots per inch.

## ACKNOWLEDGEMENTS

We thank Sarah Antinone for providing embryonic chick DRGs, Duncan Wilson and Prashant Desai for generously providing antibodies, and Ekaterina Heldwein for her assistance designing the VP5 Δ6F10 mutant. Transmission electron microscopy samples were processed by Northwestern University’s Center for Advanced Microscopy and imaged using the facility’s FEI Tecnai Spirit G2 transmission electron microscope. Sequencing services were performed at the Northwestern University Genomics Core Facility.

